# Computational analysis of the effect of SARS-CoV-2 variant Omicron Spike protein mutations on dynamics, ACE2 binding and propensity for immune escape

**DOI:** 10.1101/2021.12.14.472622

**Authors:** Natalia Teruel, Matthew Crown, Matthew Bashton, Rafael Najmanovich

## Abstract

The recently reported Omicron (B.1.1.529) SARS-CoV-2 variant has a large number of mutations in the Spike (S) protein compared to previous variants. Here we evaluate the potential effect of Omicron S mutations on S protein dynamics and ACE2 binding as contributing factors to infectivity as well as propensity for immune escape. We define a consensus set of mutations from 77 sequences assigned as Omicron in GISAID as of November 25. We create structural models of the Omicron S protein in the open and closed states, as part of a complex with ACE2 and for each of 77 complexes of S bound to different antibodies with known structures. We have previously utilized Dynamical Signatures (DS) and the Vibrational Entropy Score (VDS) to evaluate the propensity of S variants to favour the open state. Here, we introduce the Binding Influence Score (BIS) to evaluate the influence of mutations on binding affinity based on the net gain or loss of interactions within the protein-protein interface. BIS shows excellent correlation with experimental data (Pearson correlation coefficient of 0.87) on individual mutations in the ACE2 interface for the Alpha, Beta, Gamma, Delta and Omicron variants combined. On the one hand, the DS of Omicron highly favours a more rigid open state and a more flexible closed state with the largest VDS of all variants to date, suggesting a large increase in the chances to interact with ACE2. On the other hand, the BIS shows that apart from N501Y, all other mutations in the interface reduce ACE2 binding affinity. VDS and BIS show opposing effects on the overall effectiveness of Omicron mutations to promote binding to ACE2 and therefore initiate infection. To evaluate the propensity for immune escape we calculated the net change of favourable and unfavourable interactions within each S-antibody interface. The net change of interactions shows a positive score (a net increase of favourable interactions and decrease of unfavourable ones) for 41 out of 77 antibodies, a nil score for 15 and a negative score for 21 antibodies. Therefore, in only 28% of S-antibody complexes (21/77) we predict some level of immune escape due to a weakening of the interactions with Omicron S. Considering that most antibody epitopes and the mutations are within the S-ACE2 interface our results suggest that mutations within the RBD of Omicron may give rise to only partial immune escape, which comes at the expense of reduced ACE2 binding affinity. However, this reduced ACE2 affinity appears to have been offset by increasing the occupancy of the open state of the Spike protein.

## 1. Introduction

Early samples of the Omicron variant of concern of SARS-CoV-2 were first sequenced by the Botswana Harvard HIV Reference Laboratory in South Africa on the 22^nd^ November 2021 (GISAID: EPI_ISL_6752027). Samples from Gaborone showed a cluster of sequences with a high number of variants, in total 60 such mutations when compared to Wuhan-Hu-1 (RefSeq NC_045512.2), and 50 nonsynonymous variants that changed amino acid sequence of viral proteins. Of particular concern was the presence of 32 of these in the Spike protein which interacts with ACE2 to gain entry into human cells. Of these, 15 occur within the receptor binding domain (RBD) which is both critical to ACE2 interaction and a site which overlaps with the majority of antibody epitopes [1,2], raising widespread concerns of immune escape. Specifically, of the 32 Spike protein amino acid variants in Omicron, 6 are involved in the hydrogen bonding network with, and make direct contacts to, ACE2. A further 12 residues that have changed contribute to Barnes epitope classes 1, 2 and 3 [1,2]. Comparing to previous WHO variants of concern: Alpha contains 8 Spike AA mutations, with only N501Y participating in ACE2 interaction and Barnes epitope classes 1 and 2. Beta contains 10 Spike mutations, of which 2 participate in ACE2 contact, and 3 contribute to Barnes epitope classes 1, 2 and 3. Additionally, in Beta, 2 mutated residues occur in antigenic supersite i [3]. Gamma contains 12 Spike mutations, with 2 participating in hydrogen bonding and contacts with ACE2, and 3 residues which are within Barnes epitope classes 1-3. Similarly to Beta, Gamma also contains 2 mutated residues which are within antigenic supersite i. The current dominant variant, Delta, possesses 9 Spike mutations, 4 of which are within antigenic supersite i and 1 of which is in Barnes epitope class 2. It should be noted that Alpha, Delta and Omicron possess similar mutations at the furin cleavage site (P681H, P681H and P681R respectively) outside the RBD, which show higher fusogenic potential, potentially increasing infectivity [3–5]. On the basis of the sheer number of changes in Omicron that affect both ACE2 interactions and known epitope classes as well as the addition of enhanced furin cleavage, at least on paper Omicron represents a clear cause for concern with potential to outdo previous aforementioned variants of concern.

The Pango Network [6] designated these sequences first observed in South Africa as a branch of the B.1.1 lineage, specifically B.1.1.529, on Wednesday 24^th^ November 2021. The World Health Organisation (WHO) subsequently declared B.1.1.529 a variant of concern with the designation Omicron on 26^th^ November 2021. Whilst immune escape studies are actively being undertaken by pharma, WHO and research labs alike, results are still not fully known, although early indications are some degree of immune escape is present [https://assets.publishing.service.gov.uk/government/uploads/system/uploads/attachment_data/file/1040076/Technical_Briefing_31.pdf]. We use computational structural biology methods to investigate the Spike protein mutations to evaluate the key concerns of infectivity and immune escape properties of Omicron. Our methodology can be applied to any combination of Spike protein variants, it builds on and aids our understanding of how individual amino acid changes contribute to both immune escape and increased viral infectivity. Furthermore, precisely because protein structure can tolerate individual amino acid substitutions with small shifts and rigid-body movements [7–12], our work emphasises the need to understand the nature of the amino acid changes and the roles the residues play to provide a robust interpretation of the dangers of new variants rather than raising concerns due to total numbers of non-synonymous changes alone.

## 2. Methods Highlights

The consensus sequence of Omicron S was obtained from an initial set of 77 sequences from GISAID [13,14] available on November 25, 2021 with a 50% sequence identity threshold. The set of mutations employed here for the most part still is part of the definition of the Omicron S protein. The mutations considered are the following: A67V, T95I, G142D, L212I, G339D, S371L, S373P, S375F, K417N, N440K, G446S, S477N, T478K, E484A, Q493R, G496S, Q498R, N501Y, Y505H, T547K, D614G, H655Y, N679K, P681H, N764K, D796Y, N856K, Q954H, N969K, L981F. A number of deletions and insertions exist in the Omicron S protein but are ignored here due to the difficulty in modelling the precise effect of deletions and insertions on protein structure. The structure of the open and closed states of the trimeric S Omicron protein as well as Omicron S in complex with ACE2 and 77 antibodies was prepared in the same manner as discussed in [15].

Dynamical Signatures (DS) and Vibrational Difference Scores (VDS) are based on normal mode analysis [16] and were introduced in [15] for the study of single point mutations and previous variants of the SARS-CoV-2 S protein.

Binding residues were determined based on calculated atomic surface areas in contact [17]. The overall interaction between two residues was determined as the weighted sum of atomic surface areas in contact using a matrix of pairwise pseudo-energetic interaction scores for 40 atom types as used in the docking program FlexAID [18]. When this sum is negative, the two residues in question are said to have a favourable interaction. Contrarywise, if the sum is positive, the two residues are said to have an unfavourable interaction. Considering the limitations of the pseudo-energetic definition of interaction scores, we restrict the analysis to determining if the interaction between two residues is favourable or unfavourable. When the Omicron S protein in complex with either ACE2 or an antibody is analysed relative to the wildtype, there are favourable and unfavourable interactions gained and lost, not only directly at the mutated position but also elsewhere as the introduction of mutations affects the entire interface. Therefore, relative to the wildtype, a given position will have a net change in the number of interactions. A positive value denotes the combined effect of gaining favourable interactions and loosing unfavourable ones. Such a Net Change in Interactions (NCI) can be defined for every residue and summed for the entire interface. To understand the specific effect of a mutation, we can combine the NCI of a residue and those of its nearest neighbours in 3D as the effect of a mutation is predominantly local but can involve not only its own interactions but those of residues around it. We define a Binding Influence Score (BIS) for a given residue *i* as the sum of its BIS and those of its neighbours within 5A with a 0.1 weight factor, except for neighbour residues that are themselves mutating:

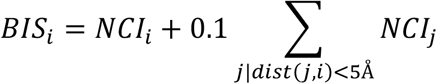

## 3. Results

Calculation of the Dynamical Signature of the Omicron S protein shows a large rigidification of the open state and some increase of flexibility of the closed state (Figure 1) with contributions from multiple residues but primarily concentrated on the receptor binding domain (RBD), leading to a Vibrational Difference Score of 7.66×10^-1^J.K^-1^. This is the largest VDS of all variants calculated thus far taking into account scores from our previous work on other variants and mutations of concern [3] (Table 1).

**Figure 1.**
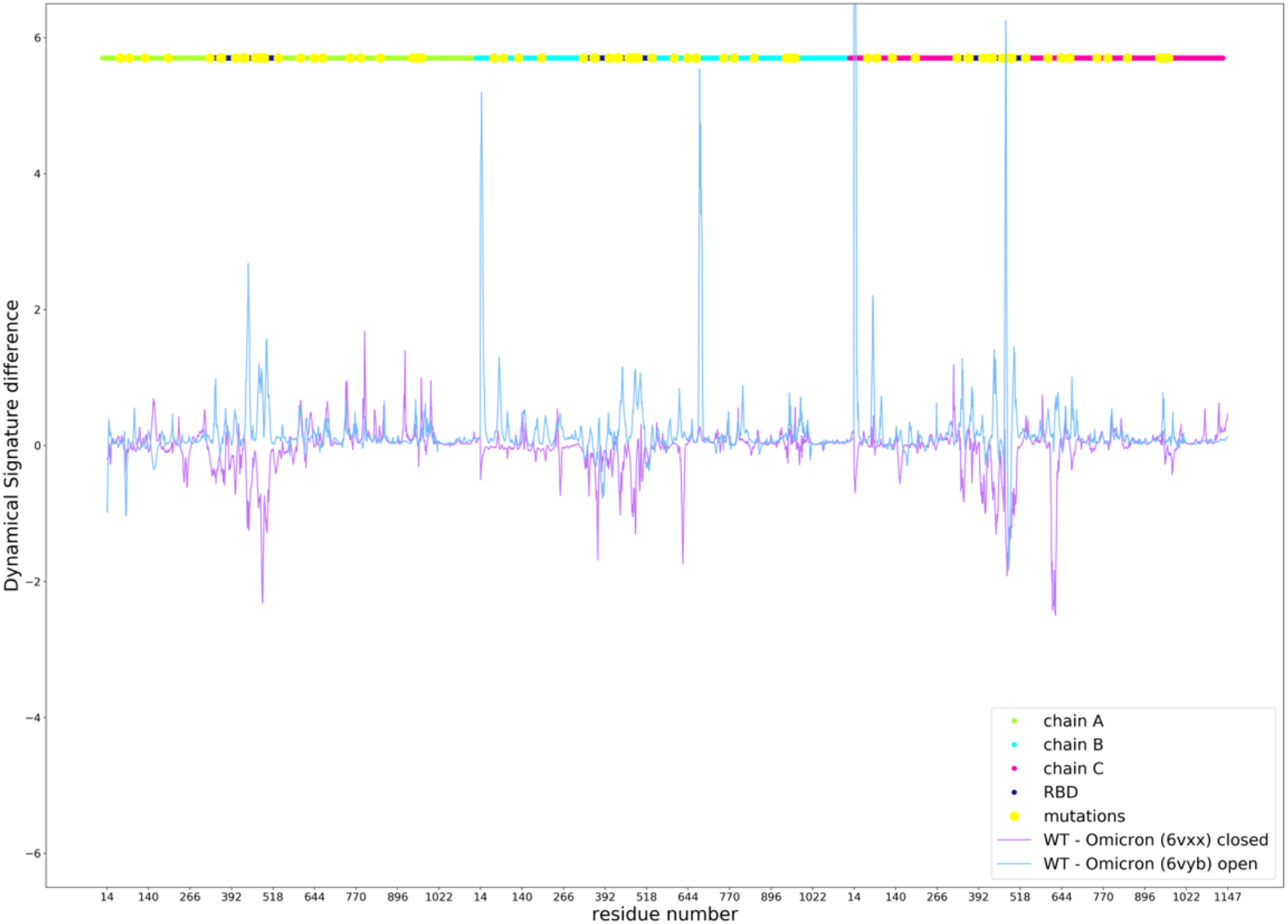
Dynamical Signature of the Omicron S protein open and closed state compared to the wildtype. Positive values indicate an increase in rigidification and negative values indicate a decrease in rigidity. X axis shows the residue number, which iterates thought all residues for chains A (green), B (cyan), and C (magenta) following on from each other. Mutations are depicted in yellow.

**Table 1.**
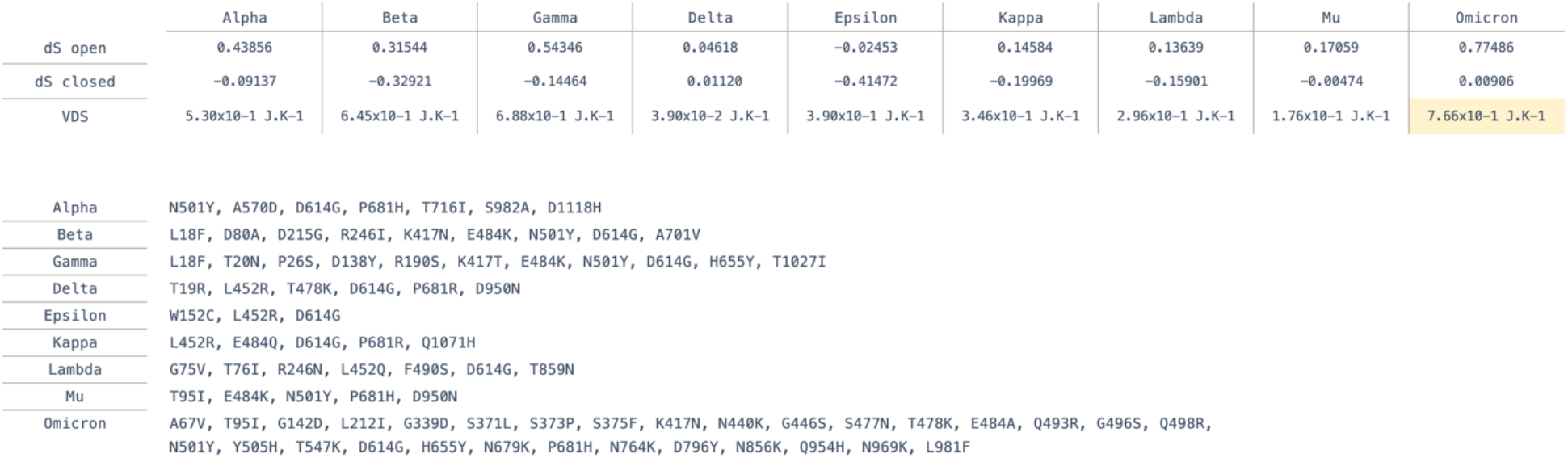
Definition of point mutations in variants and their calculated Vibrational Difference Score (VDS).

Our previous study of VDS on past variants suggests that an increase in VDS is associated with longer occupancy of the open state required for binding ACE2, higher VDS values thus favour interactions with ACE2 [15]. This could contribute to a potential higher infectivity of Omicron, as was the case for the Alpha variant that appeared earlier during the pandemic [3]. However, past variants contained fewer mutations than Omicron and were to a great extent dominated by the effect of a single mutation or a few mutations. Specifically, N501Y increases VDS and binding affinity to ACE2 as determined experimentally [19] as well as through our BIS analysis (see below) and shown to increase infection and transmission [20].

We calculate Binding Influence Scores (BIS) for the interactions between S-ACE2 interface residues for all mutated interface residues in several variants and compare these results to the experimental result by Starr *et al*. [19]. Our calculated BIS values show a Pearson correlation Coefficient of 0.87 with the experimental results (Figure 2).

**Figure 2.**
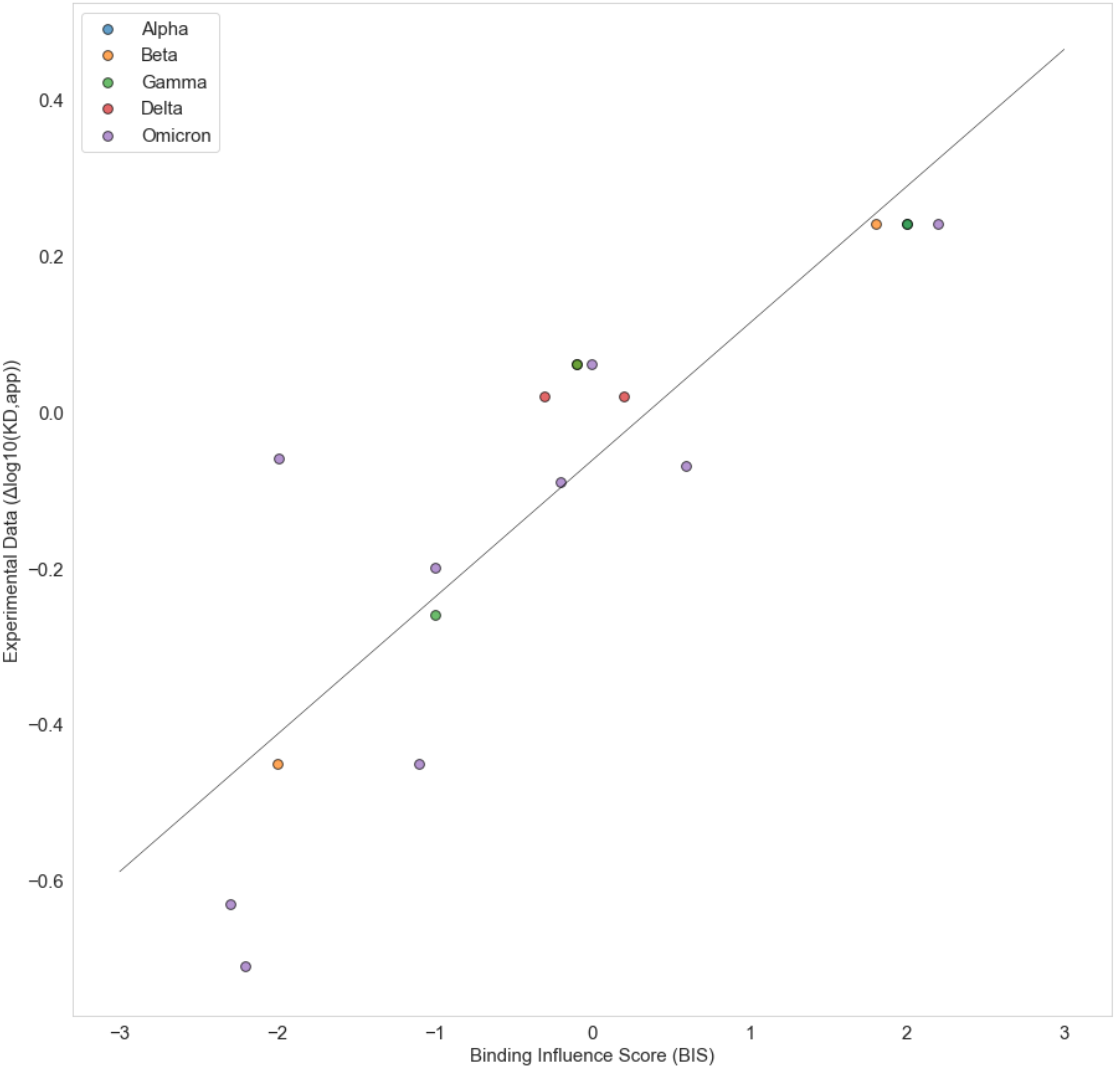
Correlation between Binding Influence Scores (BIS) and experimental values of the effect of single point mutation on Spike ACE2 binding affinity [19].

Most mutated residues in the Omicron S-ACE2 interface decrease binding affinity as observed experimentally when appearing independently but also together as in Omicron S according to our BIS calculations (Table 2).

**Table 2.**
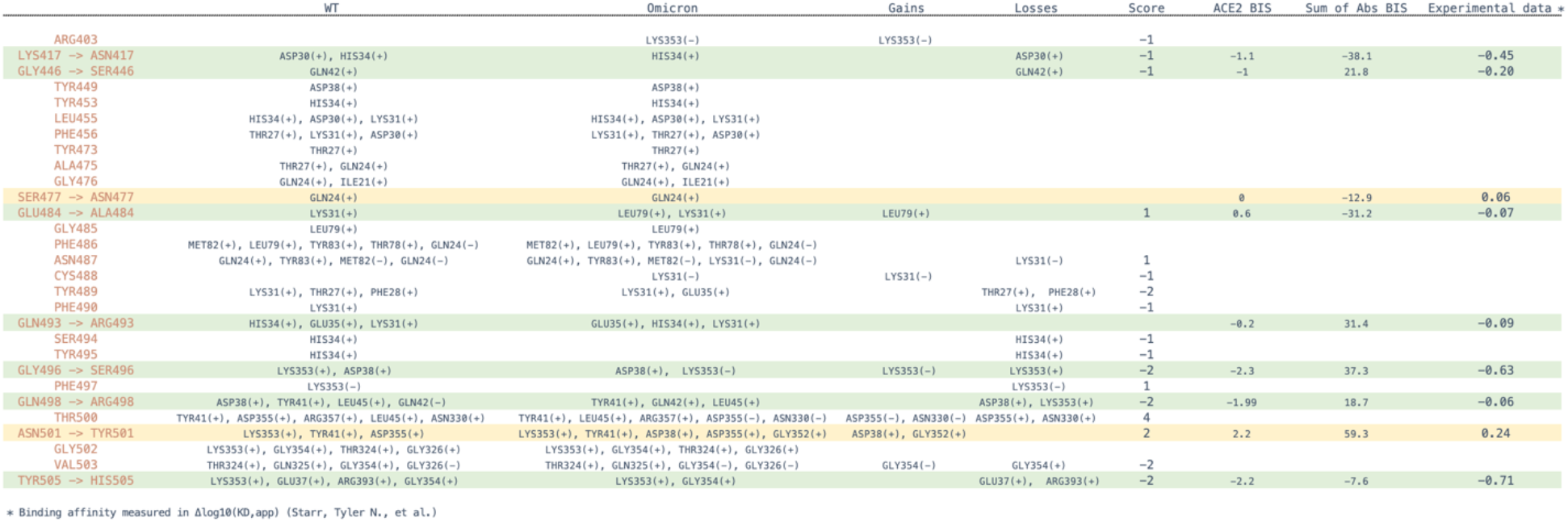
Specific changes in favorable and unfavorable interactions for Omicron S relative to the wildtype and their effect on BIS. Spike mutations in Omicron are in red for each row, interacting partner residues in ACE2 are listed in columns WT and Omicron for each variant. Binding Influence Scores (BIS) are calculated for ACE2 interactions in ACE2 BIS. Negative and positive BIS values means reduced or increased binding affinity respectively. The sum of BIS scores for 77 antibodies are given in Sum of Abs BIS column. The final column is a sum of escape scores for each mutation by Starr *et al*. [19].

Among the Omicron S-ACE2 interface mutated residues, five residues, namely positions 417, 446, 496, 501, 505 have the strongest effects on binding based on the experimental data by Starr *et al*. [19]. When calculating the sum of BIS values for the same residues across all 77 antibodies, we observe that three of those have an equivalent predicted effect on antibody binding. The majority of antibodies interact with the SARS-CoV-2 S protein through the same interface as that which is used to bind to ACE2 (Figure 3). With that in mind, our results hint at the possibility that the evolution of SARS-CoV-2 is selecting in part residues that facilitate immune escape at the expense of ACE2 binding affinity.

**Figure 3.**
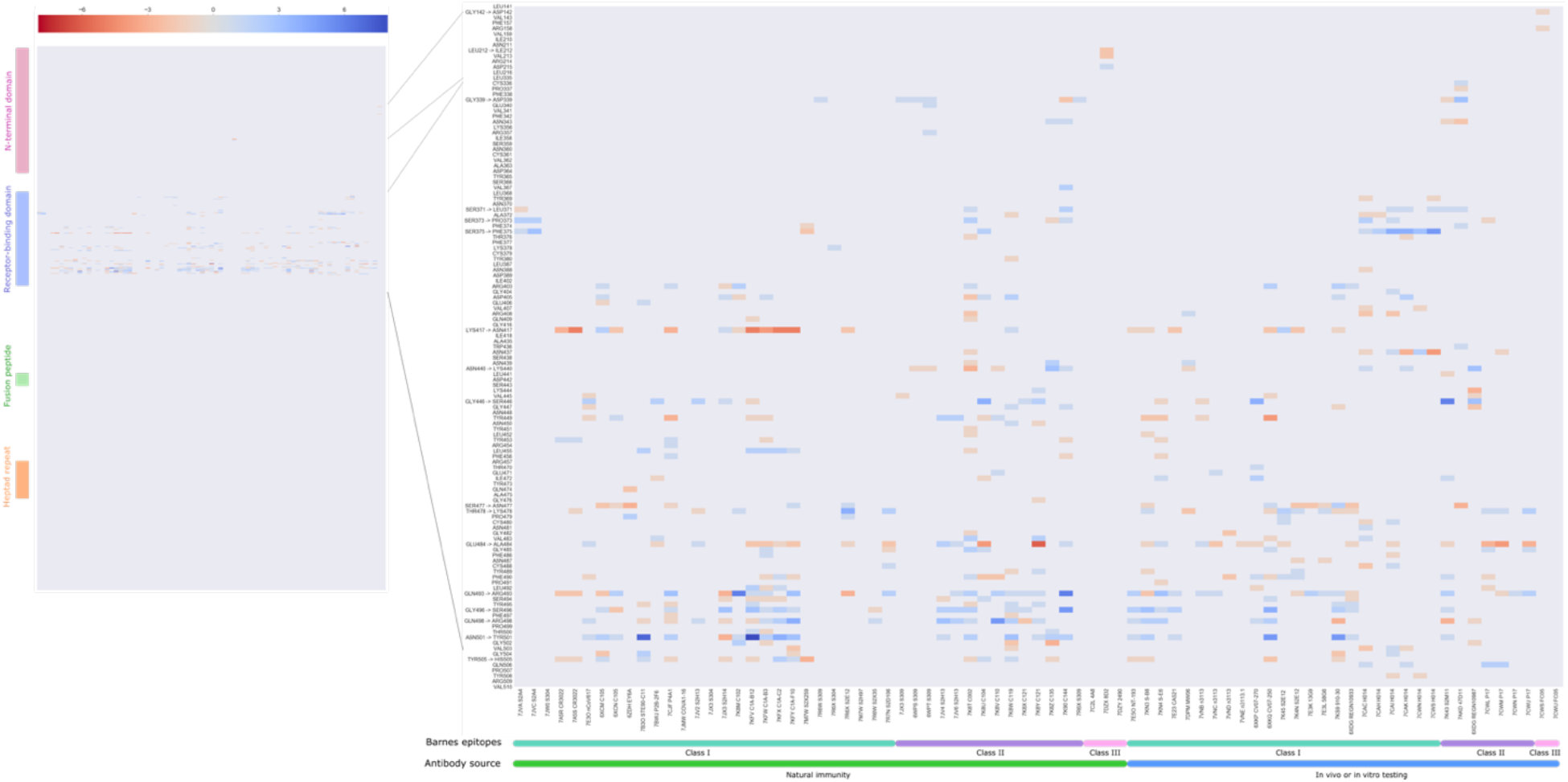
Heatmap of the total score of interactions between Spike residues (rows) and 77 antibodies from cocrystallized structures (columns). This dataset of 77 antibodies is divided according to source - infection acquired immunity or antibodies generated in animal models or tested *in vitro* – and according to the class of the epitope following Barnes *et al.* [2].

Analysis of the net change in interactions (NCI) for each antibody as a whole shows that for 41 antibodies in the dataset the overall effect is in fact one of an increase in favourable interactions, despite the fact that some important mutations by themselves lead to a decrease in affinity. Only for 21 out of the 77 antibodies, a decrease of NCI is observed (Supplementary data) as shown in Figure 4.

**Figure 4.**
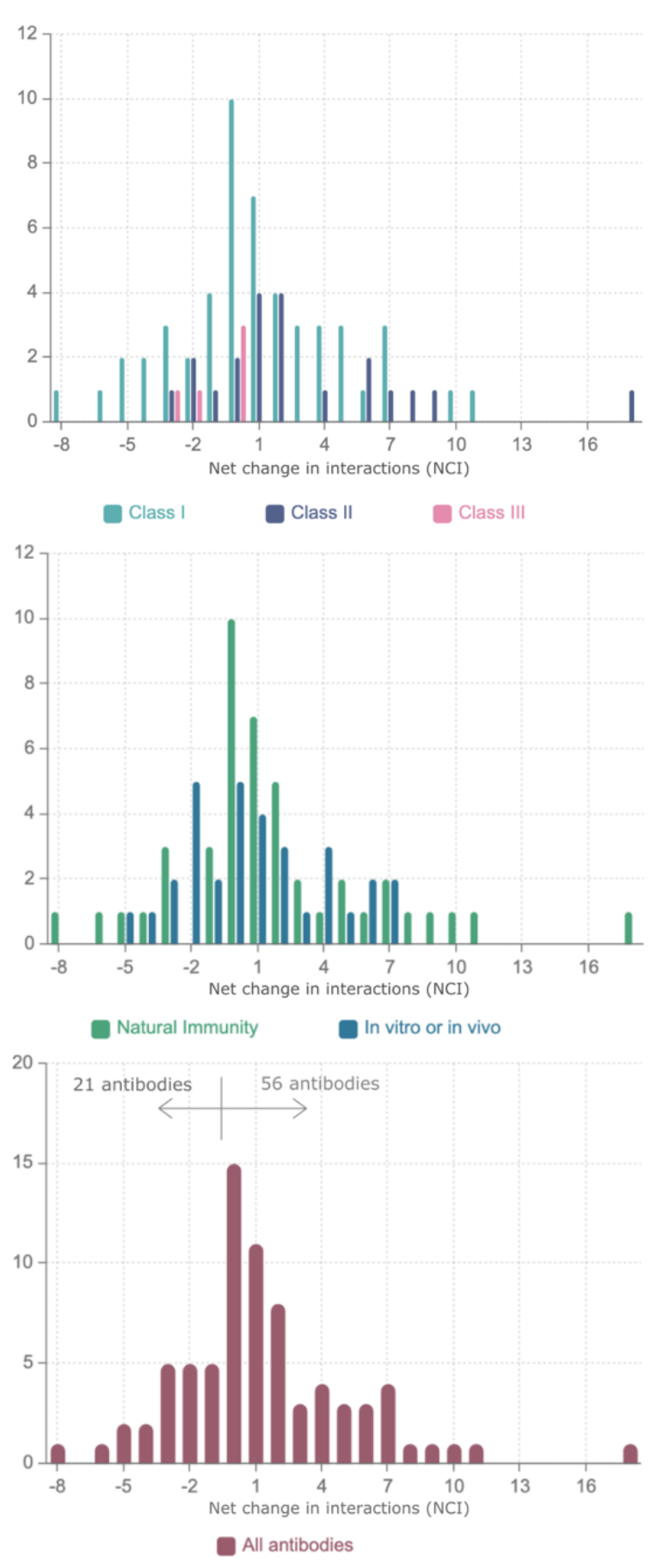
Distribution of Net Change in Interactions (NCI) for a dataset of 77 antibodies divided according to the class of the epitope according to Barnes *et al.* [2] and representing infection acquired immunity as well antibodies generated in animal models or tested *in vitro*.

Lastly, Figure 5 shows a summary of our findings, depicting from left to right panels, the position of Omicron mutations, their overall effect on dynamics and their potential in facilitating immune escape.

**Figure 5.**
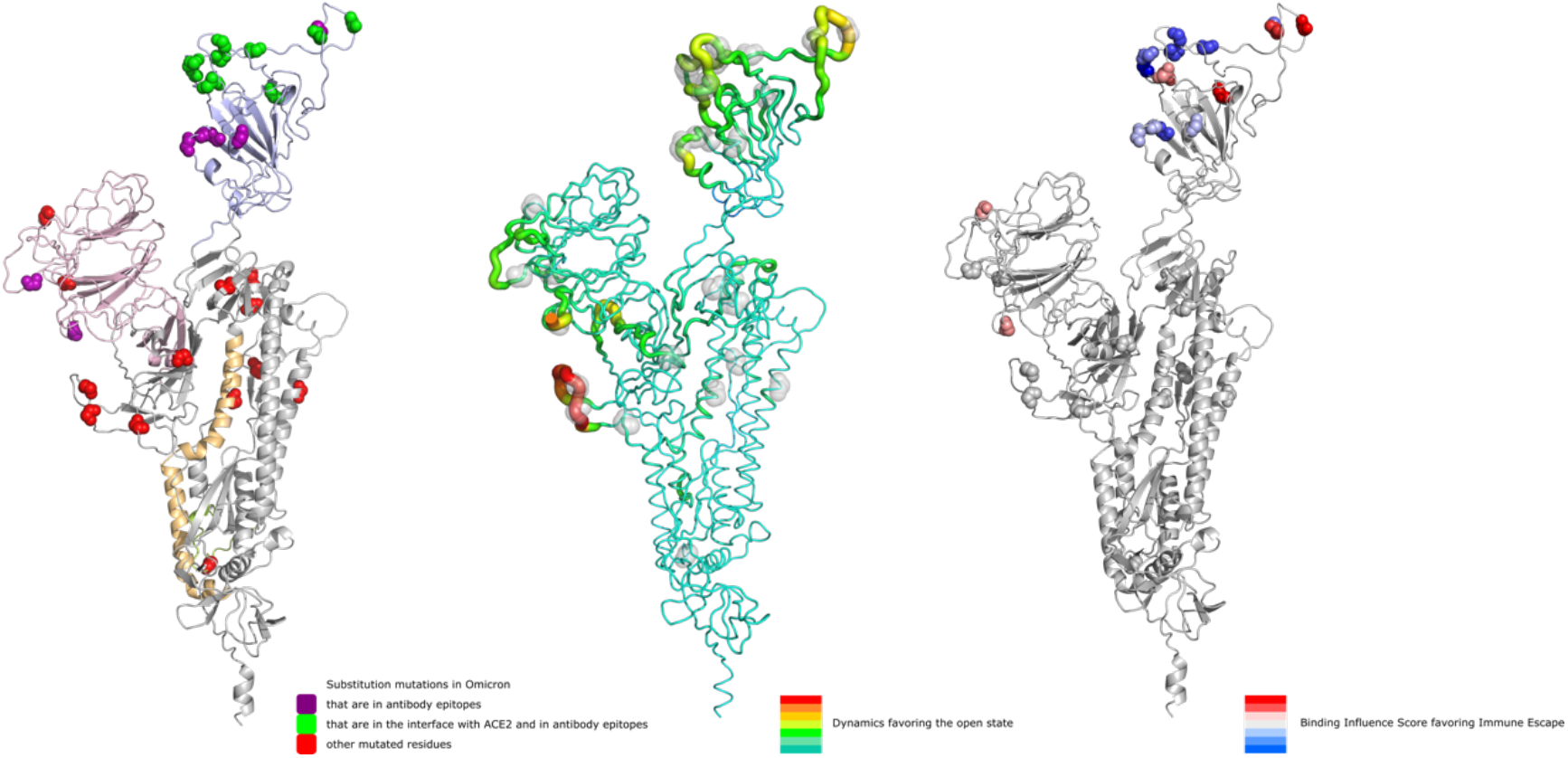
Depiction of substitution mutations in Omicron, their effect on dynamics favoring the open state (increasing from green to red) and their potential for immune escape (increasing from blue to red).

### Data availability

The data generated in this study and supplementary file can be accessed at the following Git depository: https://github.com/nataliateruel/Omicron_data

## Conclusions

The mutations in the S protein of the Omicron variant seem to facilitate interactions with ACE2 through a decrease of flexibility of the open state and concomitant increase in flexibility of the closed state. At the same time, most of the mutations in the Omicron S-ACE2 interface seem to decrease ACE2 interaction affinity. This is possibly arising from selection pressure to facilitate immune escape as a considerable number of antibodies target the same interface. However, our calculations suggest that immune escape is only partial and in fact interactions with several antibodies may become stronger. It is unclear to what extent the antibodies in our dataset are representative of what different vaccines may elicit. Assuming that they are representative, we believe that the Omicron variant poses only a partial immune escape risk to individuals exhibiting a strong immune response to SARS-CoV-2 developed prior to the appearance of the Omicron variant.

A recent preprint [21] utilized AlphaFold [22] to model the Omicron S protein and the proteinprotein docking software Haddock [23] to predict S-Antibody interactions for four antibodies. It is unclear how small variations in local structure of S as modelled by AlphaFold or of S-antibody complexes as a result of the docking simulation may affect the results. The method presented here on the other hand, ignores deletions and insertions but uses the experimentally determined S-antibody structures instead of docking poses for a considerably larger set of antibodies. Whereas previous variants contained fewer mutations in the S protein and with clear effects on both binding and dynamics (such as N501Y containing variants), Omicron has a larger set of mutations (with more potential for errors in modelling for example) that brings about more mixed results (larger VDS but weaker predicted binding). As the results obtained by Ford *et al*. [21] and those presented here are in agreement, we believe that ignoring insertions and deletions does not substantially impact our results. Considering the fast execution time for our BIS analysis, which makes it possible to analyse large datasets of antibodies, we plan to increase this dataset as more S-antibody structures become available to monitor DS, VDS and BIS for future variants of concern.

Our work highlights the potential evolutionary trade-off between ACE2 binding and immune escape operating as a joint selection pressure on the RBD of the spike protein, it is likely that Omicron has managed to balance the reduced S-ACE2 affinity by compensatory mutations that stabilise the open state of the S protein. Another important point to emphasize is that these open conformation stabilizing mutations (or closed state destabilization) are not exclusively located in the RBD and this demonstrates the importance of understanding the far-reaching effect of mutations on the spike protein through an understanding of the dynamics of the protein structure.

We believe that the Omicron SARS-CoV-2 variant is more transmissible given the very high VDS values compared to other variants and can evade immune response to some extent but is not likely to pose a serious personal risk to individuals with a strong immune response to past variants given the weaker binding to ACE2 and stronger binding to the majority of antibodies tested here.

## Acknowledgments

MB holds COG-UK funding and is currently supporting SARS-CoV-2 sequencing funded by the UK Health Security Agency. MC is supported in this work by COG-UK funding. Both MB and MC are supported by Research England’s Expanding Excellence in England (E3) fund. RN is a Fonds de Recherche du Québec - Santé (FRQ-S) Senior Fellow, a member of the Réseau Québécois de Recheche sur les Médicaments (RQRM) and the Quebec Network for Research on Protein Function, Engineering and Applications (PROTEO). This work was funded by grants from Genome Canada (http://genomecanada.ca), Genome Québec (https://www.genomequebec.com) as well as the Natural Sciences and Engineering Research Council (NSERC) grant number RGPIN05332-2019. This research was enabled in part by support provided by (Calcul Québec) (https://www.calculquebec.ca/) and Compute Canada (www.computecanada.ca).

